# Interleukin-22 drives a metabolic adaptive reprogramming to maintain mitochondrial fitness and treat liver injury

**DOI:** 10.1101/2020.01.02.892927

**Authors:** Wei Chen, Wenjing Zai, Jiajun Fan, Xuyao Zhang, Jingyun Luan, Yichen Wang, Yilan Shen, Ziyu Wang, Shixuan Dai, Si Fang, Dianwen Ju

**Affiliations:** Minhang Branch, Zhongshan Hospital, Fudan University/Institute of Fudan-Minhang Academic Health System, Minhang Hospital, Fudan University; Department of Biological Medicines, Fudan University School of Pharmacy, 826 Zhangheng Road, Shanghai, 201203, P. R. China; Department of Ophthalmology, Stanford University School of Medicine, Palo Alto CA 94304, USA; Key Laboratory of Medical Molecular Virology, School of Basic Medical Sciences, Shanghai Medical College of Fudan University; Department of Nephrology, Changhai Hospital, Second Military Medical University, Shanghai, 200433, P. R. China; Department of Pharmacy, Huadong Hospital, Fudan University, Shanghai, 200040, P.R. China; Tongcheng Hospital of Traditional Chinese Medicine, Anhui, 231400, P. R. China

**Keywords:** oxidative phosphorylation, glycolysis, mitochondria, reactive oxygen species, lncRNA H19

## Abstract

Interleukin 22 (IL-22) is an epithelial survival cytokine that is at present being explored as therapeutic agents for acute and chronic liver injury. However, its molecular basis of protective activities remains poorly understood. Here we demonstrate that IL-22 inhibits the deteriorating metabolic states induced by stimuli in hepatocytes. Specifically, we provide evidence that IL-22 promotes oxidative phosphorylation (OXPHOS) and glycolysis and regulates the metabolic reprogramming related transcriptional responses. IL-22 controls metabolic regulators and enzymes activity through the induction of AMP-activated protein kinase (AMPK), AKT and mammalian target of rapamycin (mTOR), thereby ameliorating mitochondrial. The upstream effector lncRNA H19 also participates in the controlling of these metabolic processes in hepatocytes. Importantly, amelioration of liver injury by IL-22 through activation of metabolism relevant signaling and regulation of mitochondrial function are further demonstrated in cisplatin-induced liver injury and steatohepatitis. Collectively, our results reveal a novel mechanism underscoring the regulation of metabolic profiles of hepatocytes by IL-22 during liver injury, which might provide useful insights from the bench to the clinic in treating and preventing liver diseases.

**Graphical Abstract:** Our works demonstrate a critical role of IL-22 in regulating hepatocellular metabolism to treat liver injury via activating STAT3-lncRNA H19-AMPK-AKT-mTOR axis. These findings describe a novel mechanism underscoring the regulation of metabolic states of hepatocytes by IL-22 during liver injury with potentially broad therapeutic insights.

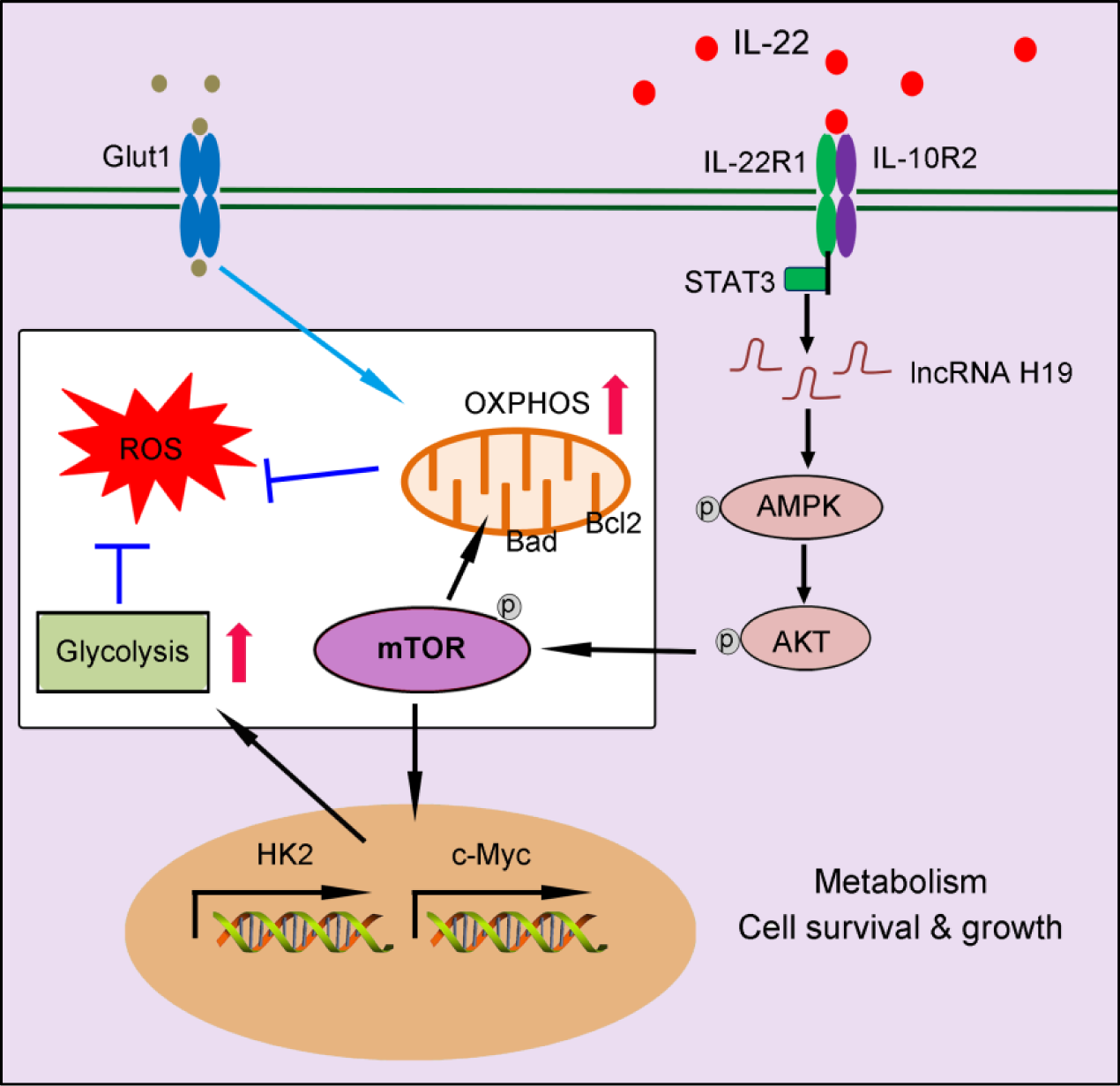

## 1. Introduction

Over the last decades, liver injury has become one of the major causes of illness and mortality with an amazing worldwide prevalence [1]. Characterized as hepatocyte damage with severe inflammation, steatosis and fibrosis, liver injury can deteriorate to end-stage cirrhosis and liver failure [2–4]. Unfortunately, evidence based-treatment for tackling liver injury is so far unavailable in clinical settings. Moreover, efforts to ameliorate the injury of hepatocytes by regulating their complications, such as intracellular oxidative stress, protein aggregates, and dysfunctional organelles, are far from satisfactory [5–8]. Thus, it is of immediate significance to explore effective pharmacological antidotes and strategies for liver injury treatment.

Interleukin-22 (IL-22), which belongs to the interleukin-10 (IL-10) cytokine family, is produced by immune cells, including T cells, NK cells, innate lymphoid cells (ILCs), as well as neutrophils, and directly regulates the function of hepatocytes [9–12]. We and others have suggested that IL-22 can protect against liver injury in several mouse models including acetaminophen, T cell or alcohol-induced hepatitis, etc. [13–17]. Despite growing evidence suggests that IL-22 is a promising antidote to treat various hepatic disorders, the molecular basis of the liver protective activities of IL-22 needs to be fully characterized. Uncovering the underlying mechanisms of IL-22 is necessary both for understanding how IL-22 acts to prevent liver injury and for identifying critical processes as well as molecular regulators involved in addressing liver injury [18–19].

Altering cellular metabolism by specific proteins or genes can promote mammalian cell and organ repair, suggesting that the metabolic processes during tissue injury are the pivotal component of cellular functions and survive [20–22]. Moreover, these works also show that specific inhibition of oxidative phosphorylation (OXPHOS) or glycolysis negates tissue restoring beneficial effects on tissue repair. Our prior researches linking IL-22 to mitochondrial function and activation of STAT3 signaling transduction lead us to inquire whether reprogramming the metabolism of hepatocytes with IL-22 can affect its hepatoprotective capacities. Thus, we hypothesize that IL-22 could beneficially regulate liver homeostasis and hepatocyte function via regulation of metabolic states in stress environmental situations.

In the present work, we first reported that the protective functions of IL-22 were mediated by metabolic reprogramming of hepatocytes. Utilizing cell biological, molecular, and biochemical approaches, we demonstrated that IL-22 opposed the decreased OXPHOS and glycolysis in the damaged hepatocytes induced by injurious stimuli. We further indicated that these metabolic processes were associated with the activation of metabolic signaling pathways, and thereby inhibiting the development of hepatocyte injury possibly via the prevention of mitochondrial dysfunction. Moreover, we investigated the essential roles of STAT3, AMP-activated protein kinase (AMPK), AKT, mammalian target of rapamycin (mTOR) and lncRNA H19 in these metabolic processes in hepatocytes with IL-22. Altogether, these observations highlight the significance of regulating metabolic profiles in hepatocytes for treating and preventing liver injury.

## 2. Results

### IL-22 regulates mitochondrial function and glycolysis in hepatocytes on injury factors stimulation

We investigated hepatocytes, for changes in the oxygen consumption rate (OCR), and extracellular acid rate (ECAR), as a measure of OXPHOS and glycolysis, respectively. The damaged hepatocytes, which were stimulated with liver injury factors, became less oxidative and glycolysis, as shown had lower basal OCR and ECAR values (Fig. 1A and 1B). It was noteworthy that IL-22 promoted OXPHOS and glycolysis in these hepatocytes, whereas the metabolic reprogramming effects were completely disarmed by a neutralizing antibody against the IL-22 receptor (IL-22R1) indicating IL-22 promoted OXPHOS and glycolysis via targeting hepatocytes directly. (**Fig. S1**). The effects of IL-22 on mitochondrial and glycolytic flux in hepatocytes were further assessed (Fig. 1D and 1E). Just as anticipated, IL-22 reversed the stimuli-induced impairments in maximal respiratory capacity (MRC) and glycolytic flux (Fig. 1D, 1E and 1F). These results were also demonstrated by an increase in glucose uptake with the addition of exogenous IL-22 (Fig. 1G).

**Figure 1.**
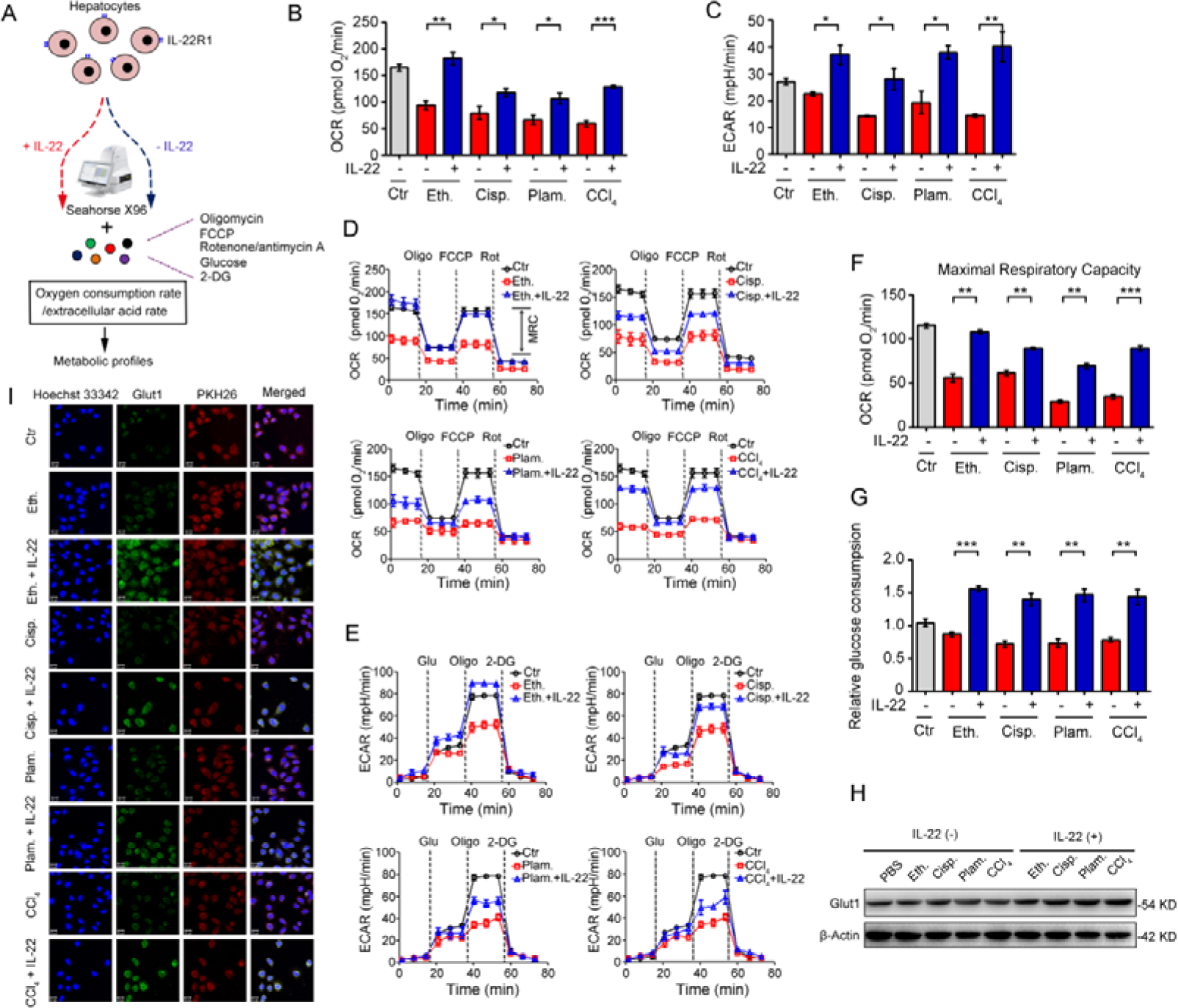
IL-22 regulates mitochondrial function and glycolysis in hepatocytes. **(A)** Using Seahorse XF96 Extracellular Flux Analyzer to assess the changes in the oxygen consumption rate and extracellular acid rate of hepatocytes. (**B** and **C**) OCR and ECAR in hepatocytes treated with 200 mM ethanol, or 5 μg/mL cisplatin, or 0.25 mM palmitic acid, or 10 mM CCl_4_ in the absence or presence of IL-22 for 24 h (*n* = 3). (**D** and **E**) Representative curves in the OCR and ECAR of hepatocytes after incubated with oligomycin, glucose, FCCP, rotenone, and 2-DG (*n* = 3). (**F**) MRC of hepatocytes evaluated by real time changes in OCR (*n* = 3). (**G**) Relative glucose consumption in hepatocytes upon IL-22 treatment (*n* = 3). (**H**) Glut1 protein expression upon IL-22 treatment under injury stress. (**I**) Localization and expression of Glut1 (green), nuclear (blue), and plasma membrane (red) in hepatocytes treated as in (**B**) for 24 h. Scale bars, 20 μm; *P < 0.05, **P < 0.01, ***P < 0.001.

Because the kinetics of glucose transporter Glut1 plays a principal role in glucose homeostasis, we therefore sought to test whether IL-22 affects Glut1 expression or translocation from intracellular to the plasma membrane in hepatocytes. The expression and localization of Glut1 were determined by western blot and the presence of co-localization between cell surface (PKH26 for cell membrane labeling, red) and Glut1 (green). Both assays indicated that IL-22 promoted the expression of Glut1 (Fig. 1H) and translocated it to the plasma membrane (Fig. 1I and **S2**). Collectively, our observations illustrated that IL-22 promoted OXPHOS and glycolysis in the damaged hepatocytes.

### IL-22 upregulates the metabolic reprogramming related transcriptional responses

To identify the transcriptional processes elicited by IL-22, we performed RNA sequencing analysis (RNA-seq) on hepatocytes by comparing injury factors-challenged groups to IL-22 plus injury factors-challenged groups at 6 h. Gene expression profiles identified 162 genes whose transcriptional levels were uniquely changed by IL-22 treatment (P < 0.01), indicating that IL-22 plays a vital role in gene expression (Fig. 2A). We next carried out Kyoto Encyclopedia of Genes and Genomes analysis (KEGG) and Gene Set Enrichment Analysis (GSEA) to recognize specific processes connected with IL-22 treatment and genotype. In the significantly upregulated transcripts, our data showed that only in IL-22-treated groups, but not control groups, an obvious enrichment of upregulated pathways associated with glycolysis, AMPK signaling pathway, nicotinate and nicotinamide metabolism, and choline metabolism in cancer was present in addition to expected processes (e.g., fat digestion and absorption, and central carbon metabolism in cancer; Fig. 2B-D and **Fig. S3**). To further explore our hypothesis, we evaluated the specifically down- and up-regulated genes and suggested a tremendous number of upregulated transcripts well-known to play an important part in metabolic pathways (Fig. 2E). Consistently, real-time PCR assay also showed that the metabolic enzyme (HK-2) involved in cellular metabolism and cell survival (Fig. 2F) was obviously upregulated by IL-22 treatment [20–22]. Therefore, we indicated that the upregulation of genes correlated to cellular metabolism in hepatocytes was associated with IL-22 regulated metabolic reprogramming.

**Figure 2.**
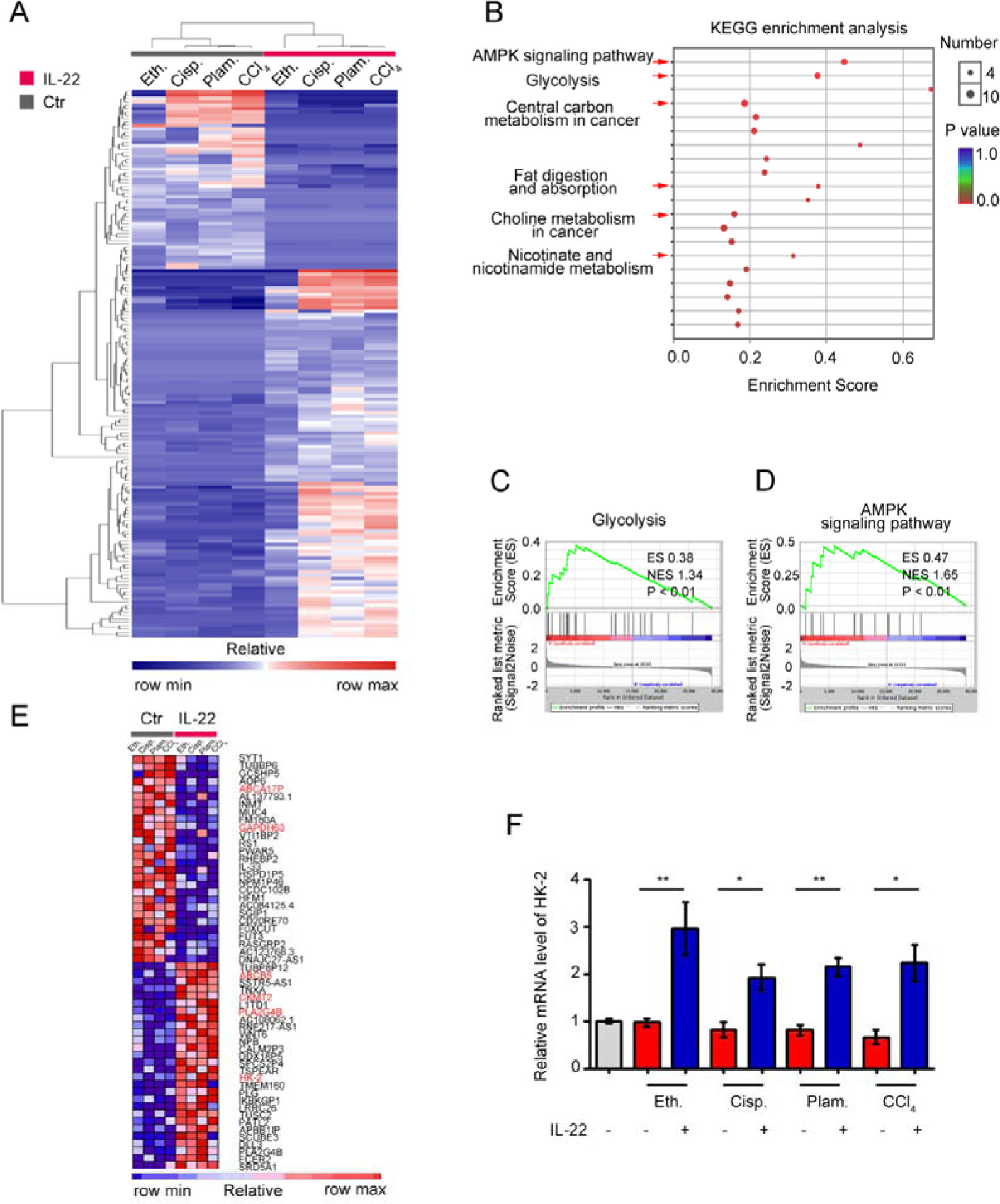
IL-22 regulates metabolic reprogramming-related transcriptional responses. (**A**) Heat map of the remarkably altered genes in hepatocytes with injury factors-challenged groups versus IL-22 plus injury factors-challenged groups. (**B**) Kyoto Encyclopedia of Genes and Genomes analysis (KEGG) of IL-22 targets in hepatocytes with IL-22-protected and -nonprotected groups. (**C** and **D**) KEGG of glycolysis and AMPK signaling pathway in IL-22-protected and -nonprotected hepatocytes. (**E**) Heat map of top altered genes from hepatocytes with IL-22 treatment. (**F**) Relative mRNA expression level of HK-2 in the absence or presence of IL-22 treatment (*n* = 3); *P < 0.05, **P < 0.01.

### IL-22 prevents the generation of dysfunctional mitochondria and mitochondrial ROS via AMPK-associated signaling mechanism

We next inquired whether the metabolic reprogramming effects of IL-22 in the damaged hepatocytes are due to the altered mitochondrial function. The hepatocytes were stained with MitoTracker Green to track the mitochondrial content. Our data suggested that the damaged hepatocytes had increased mitochondrial mass after exposure to injurious stimuli by comparison with normal hepatocytes (Fig. 3A). The phenomenon was not associated with greater hepatocytes size, because the damaged hepatocytes had similar sizes with normal cells, but rather was attributed to increased intracellular complexity indicated by the side scatter signal (SSC) (**Fig. S4)**. To differentiate between dysfunctional mitochondria and respiring mitochondria, we then stained hepatocytes with MitoTracker Red (mitochondrial membrane potential-dependent stain), and found an increase in dysfunctional mitochondria (with lower MitoTracker Red and higher MitoTracker Green) in hepatocytes after exposure to injurious stimuli (Fig. 3B). Of particular note, exogenous IL-22 was able to maintain mitochondrial fitness in damaged hepatocytes observed by decreased abnormal mitochondrial mass and dysfunctional mitochondria (Fig. 3A and 3B).

**Figure 3.**
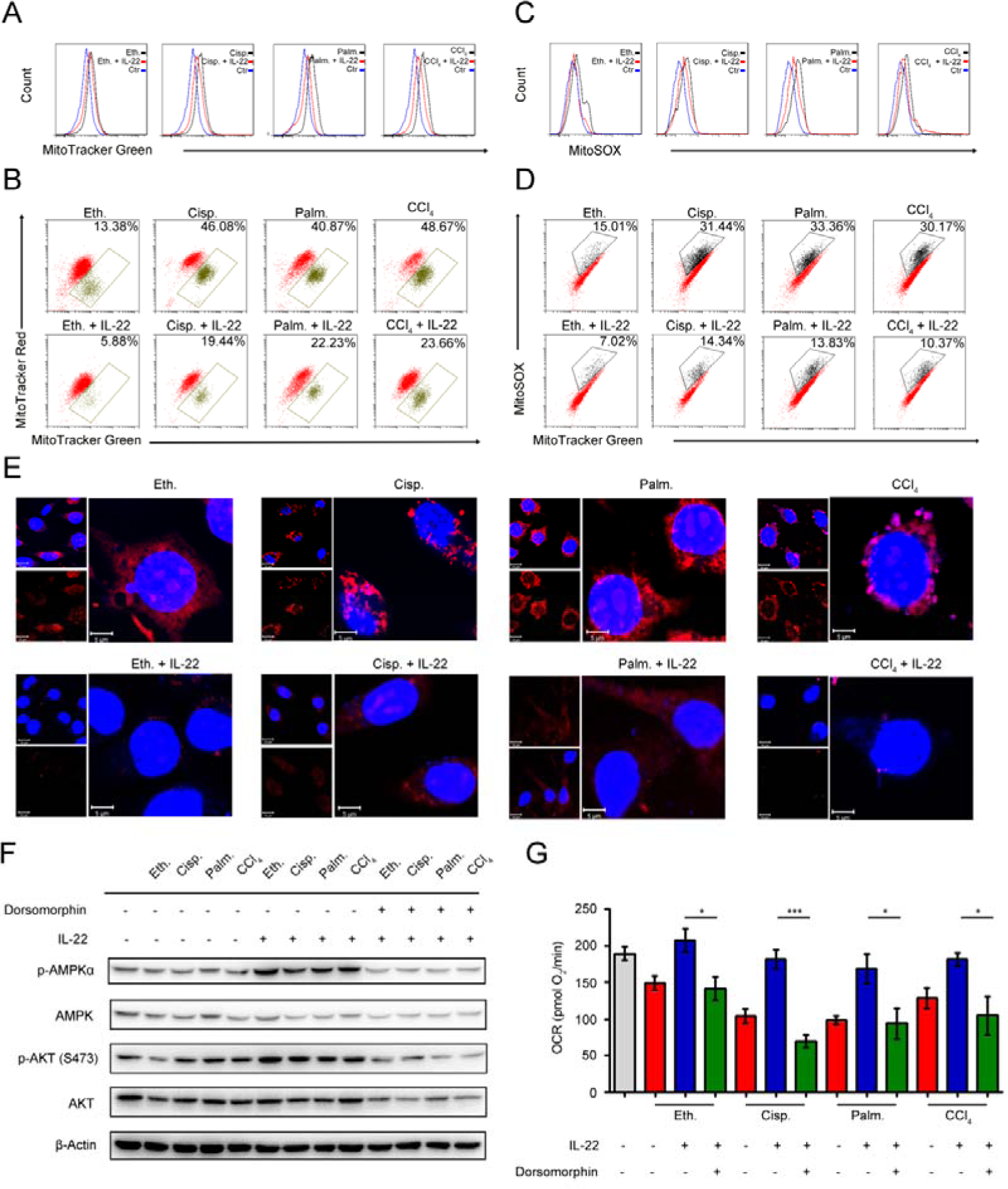
IL-22 prevents generation of dysfunctional mitochondria and mitochondrial ROS via AMPK-associated signal mechanism. Hepatocytes were treated with 200 mM ethanol, or 5 μg/mL cisplatin, or 0.25 mM palmitic acid, or 10 mM CCl_4_ in the absence or presence of IL-22 for 24 h (*n* = 3) (**A**) Mitochondrial mass was stained with MitoTracker Green and assessed by flow cytometry. Mitochondrial ROS and membrane potential were assessed in hepatocytes stained with MitoSOX (**C**), MitoTracker Green and MitoTracker Red (**B**), or MitoTracker Green and MitoSOX (**D**), respectively. (**E**) Confocal images indicated mitochondrial ROS production in hepatocytes stained with MitoSOX. (**F**) Western blot analysis suggested that IL-22 induced AMPK/AKT activation in hepatocytes, which could be inhibited by Dorsomorphin. (**G**) OCR in hepatocytes was measured in the absence or presence of indicated inhibitors and IL-22 for 24 h. *P < 0.05, **P < 0.01, ***P < 0.001.

Loss of mitochondrial membrane potential is well known to be connected with the mitochondrial ROS production [23]. We therefore asked whether the IL-22 mediated mitochondrial protective effects in damaged hepatocytes were associated with inhibition of mitochondrial ROS. As evaluated by mitochondria-specific ROS dye MitoSOX, we found that exogenous IL-22 potently inhibited the increased MitoSOX signal, which was correlated with mitochondrial content, indicating IL-22 repressed the ROS generation from dysfunctional mitochondria (Fig. 3C and 3D). These results corresponded to the live-hepatocytes fluorescence imaging, wherein the ROS accumulation could be blocked by IL-22 (Fig. 3E).

AMPK plays a key role in the cellular metabolic process via promoting catabolism to restore metabolic homeostasis, for instance, OXPHOS, glycolysis, fatty acid uptake and glucose uptake [24–25]. We observed that exogenous IL-22 induced AMPK/AKT activation in hepatocytes, suggesting that the effect of IL-22 on mitochondrial functions might AMPK/AKT associated signaling mechanism (Fig. 3F). In support of this idea, we pretreated hepatocytes with dorsomorphin (an AMPK inhibitor). As expected, the addition of IL-22 failed to maintain mitochondrial fitness (Fig. 3G). Altogether, these findings indicated that the cytokine IL-22 prevented dysfunctional mitochondria in hepatocytes via AMPK-associated signaling mechanism.

### IL-22 maintains mitochondrial function and integrity through activation of mTOR signaling transduction

mTOR is the crucial signaling hub connecting cell growth and survival, and the activation of mTOR regulates lipid synthesis, mitochondrial function and glucose metabolism [26–27]. According to the observed effects of IL-22 on mitochondrial function and cell metabolism, we tested whether IL-22 controls the activity of mTOR signaling transduction. In support of a view that mTOR might regulate metabolic homeostasis during IL-22 treatment, the addition of exogenous IL-22 to the damaged hepatocytes led to mTOR signaling transduction activation, as evidenced by increased phosphorylation of the upstream and downstream proteins such as STAT3, PI3K, S6K, S6 (Fig. 4A). Moreover, the activation of mTOR signaling transduction was inhibited in hepatocytes lacking STAT3, which suggested that IL-22 activated mTOR signaling pathway via STAT3 (Fig. 4B).

**Figure 4.**
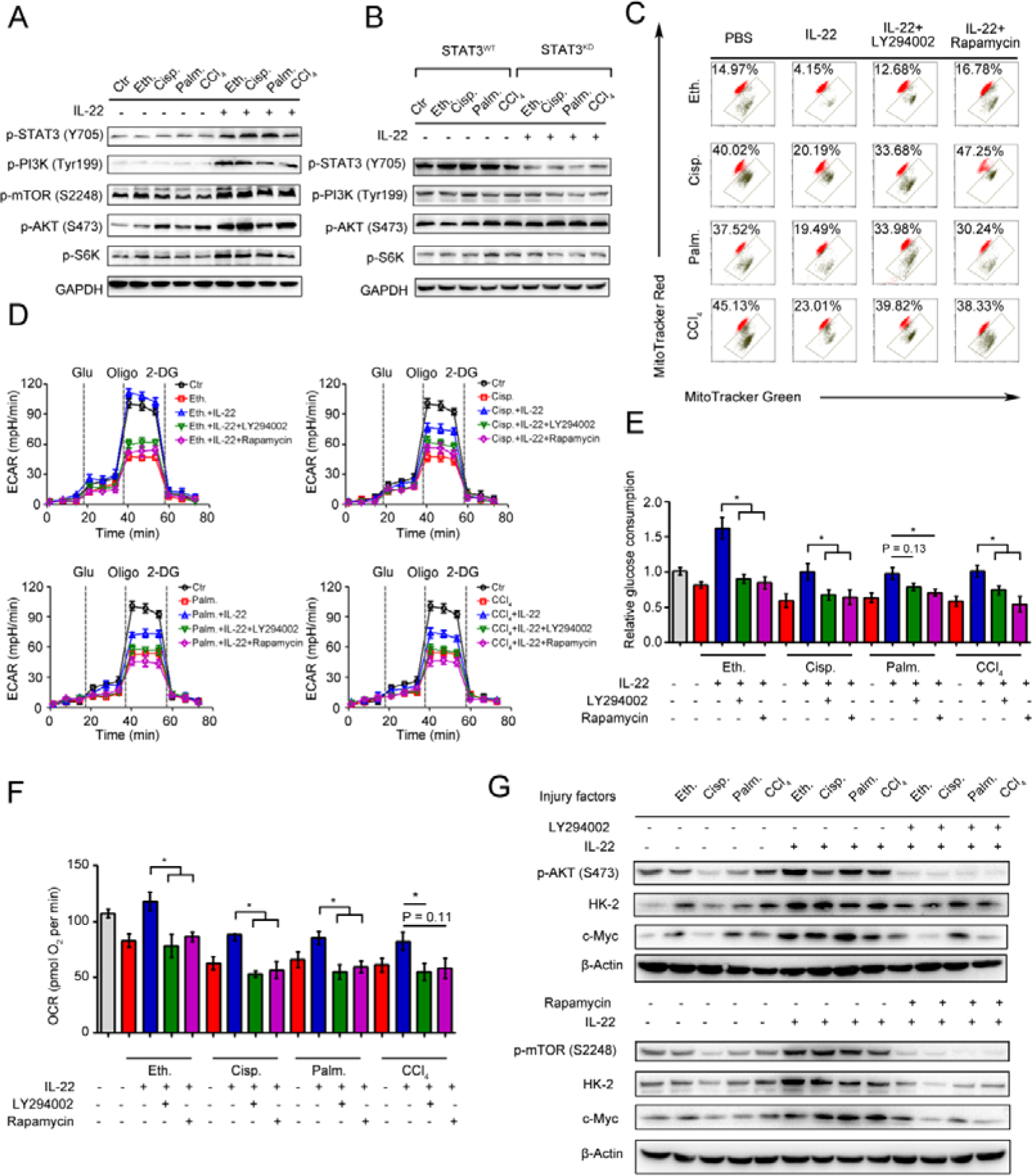
IL-22 maintains mitochondrial fitness through activation of mTOR signaling. Hepatocytes were stimulated with or without (control) 200 mM ethanol, or 5 μg/mL cisplatin, or 0.25 mM palmitic acid, or 10 mM CCl_4_ in the absence or presence of IL-22, rapamycin, or LY294002 for indicated times (*n* = 3). (**A** and **B**) Comparison of mTOR signaling activation in hepatocytes or STAT3-WT and STAT3– KD hepatocytes was assessed by western blot analysis. (**C**) Mitochondrial membrane potentials were assessed in hepatocytes stained with MitoTracker Red and MitoTracker Green. (**D**) Real time changes in the ECAR of hepatocytes after incubating with glucose, oligomycin, and 2-DG (*n* = 3) were analyzed. (**E**) Relative glucose consumption was measured for LY294002 or rapamycin treated hepatocytes at 24 hours. (**F**) Basal respiration capacity (OXPHOS) of hepatocytes was measured. (**G**) Comparison of glycolytic enzymes expression in hepatocytes versus LY294002 and rapamycin treated hepatocytes. *P < 0.05, **P < 0.01, ***P < 0.001.

We next asked whether the activation of mTOR signaling transduction by IL-22 was attributed to maintaining mitochondrial function and integrity during injury factors stimulation, which could cause mitochondrial dysfunction. We incubated hepatocytes with LY294002 and rapamycin to directly blocked mTOR signaling pathways during IL-22 treatment and investigated their oxygen consumption and mitochondrial function. Surprisingly, LY294002 and rapamycin inhibited the improved mitochondrial fitness induced by IL-22 treatment (Fig. 4C). Furthermore, co-treatment with LY294002 or rapamycin also block enhanced glycolytic flux, glucose consumption, OXPHOS and glycolytic enzymes expression, which indicated that IL-22 promoted hepatocellular metabolism through activating mTOR signaling transduction (Fig. 4D, 4E, 4F and 4G). Taken together, these results suggested that activation of mTOR signaling transduction by IL-22 treatment could result in maintained mitochondrial function and integrity.

### The induction of lncRNA H19 by IL-22 activates mTOR signaling transduction

We next sought to decipher how IL-22 activates mTOR signaling transduction. Because the activation is STAT3 dependent, the underlying mechanisms should demand transcription. To test our hypothesis, we performed gene expression analysis in hepatocytes using RNA-seq and assessed if IL-22 transcriptionally controls metabolic reprogramming, which might attribute to the activation of mTOR signaling transduction. We observed that IL-22 regulated a large number of genes, which participated in varying biological processes. Of note, from among the lncRNAs, lncRNA H19 was remarkably induced by IL-22 during injury factors stimulation (Fig. 5A). This result was also demonstrated by *in situ* hybridization assays and quantitative real-time PCR (Fig. 5B and 5C), and it required STAT3. Previous study demonstrated IL-22 activates expression of lncRNA H19 in intestinal epithelial cells (IECs), which is required for IECs healing and regeneration [28]. In the current study, we hypothesize that IL-22 drives metabolic reprogramming to maintain mitochondrial fitness and treat liver injury through lncRNA H19. To further explore the key roles of lncRNA H19 in hepatocyte metabolic states, we carried out the following series of experiments. Firstly, we found that IL-22 failed to activate AMPK/AKT/mTOR signaling transduction in hepatocytes lacking lncRNA H19 (**Fig. S5**), indicating that the activation of mTOR signaling by IL-22 was lncRNA H19-dependent (Fig. 5D and **S5**). Secondly, we observed that genetic silencing of lncRNA H19 attenuated the protective effects of IL-22 on restoring the changes in the profiles for OCR and ECAR (Fig. 5E and 5F) and the accumulation of dysfunctional mitochondria with loss of mitochondrial membrane potential and increased ROS generation after exposure to injurious stimuli (Fig. 5G and 5H), suggesting lncRNA H19 is a key target of IL-22. These data collectively suggested that the activation of mTOR signaling transduction via lncRNA H19 plays a critical role in dysfunctional mitochondria elimination in hepatocytes after exposure to injurious stimuli.

**Figure 5.**
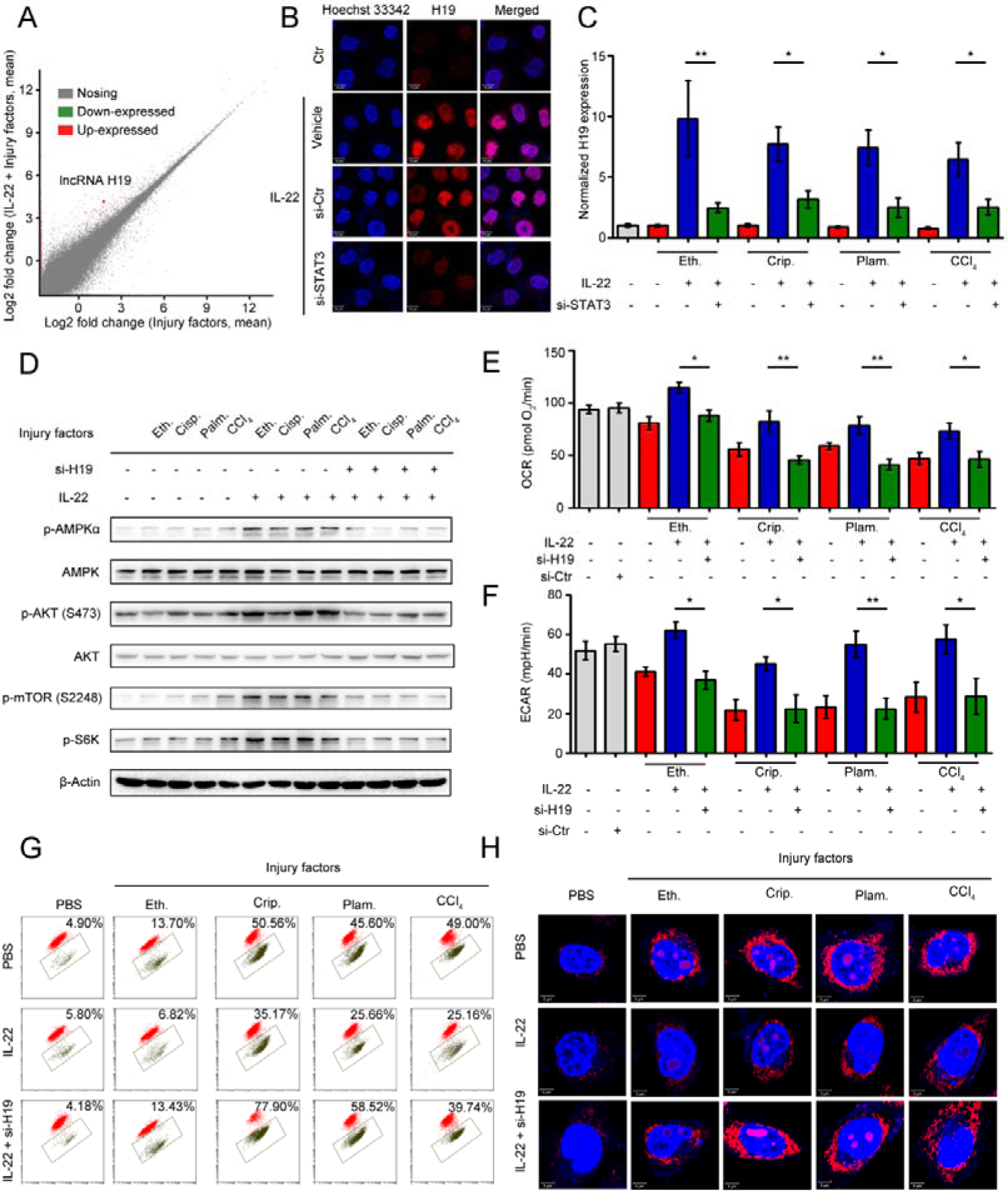
LncRNA H19 mediates the links between IL-22/IL-22R1 and the AMPK/AKT/mTOR signaling pathways. (**A**) Bland-Altman plot illustrating relative gene expression in hepatocytes stimulated with or without (control) ethanol, or cisplatin, or palmitic acid, or CCl_4_ in the absence or presence of IL-22 for the indicated times. (**B**) Confocal images suggested lncRNA H19 overexpression in hepatocytes after IL-22 treatment, which required STAT3. (**C**) Real time PCR analysis suggesting that IL-22 induced lncRNA H19 overexpression in hepatocytes, which could be prevented by STAT3 knockdown (*n* = 3). (**D**) Comparison of AMPK/AKT/mTOR signaling activation in hepatocytes versus si-H19 treated hepatocytes (lncRNA H19 knockdown). (**E** and **F**) Basal OCR and ECAR in hepatocytes at the absence or presence of IL-22, or si-H19 for 24 h (*n* = 3). (**G**) Dysfunctional mitochondria were assessed in hepatocytes stained with MitoTracker Green and MitoTracker Red. (**H**) Confocal images suggested mitochondrial ROS production in hepatocytes stained with MitoSOX.

### IL-22 attenuates hepatic oxidative stress, mitochondrial dysfunction and damage *in vivo*

Previously, we have suggested that IL-22 exerts protective potency by inhibiting ROS accumulation and preventing mitochondrial dysfunction [14–15]. To study the present model *in vivo*, we further explored mitochondrial fitness and mTOR signaling transduction in cisplatin-induced liver injury. Similar to injury factors-stimulated hepatocytes *in vitro*, animals challenged by cisplatin experienced various symptoms of hepatic injury, including hepatocyte slice degeneration and necrosis (Fig. 6B), elevation of alanine aminotransferase (ALT) and aspartate amino-transferase (AST) levels (Fig. 6C), weight reduction of livers and spleens (Fig. 6D), accumulated mitochondria with loss of mitochondrial membrane potential (MMP) (JC-1 green monomer, Fig. 6E), and increased ROS levels (Fig. 6F). However, these effects were ameliorated in IL-22-treated mice. Mechanistically, we found that IL-22 upregulated the phosphorylation of AMPK/AKT/mTOR signaling pathways as well as its downstream protein Bcl-2 (Fig. 6H). Meanwhile, Ki67, a marker of cellular proliferation, was downregulated by cisplatin stimulation, whereas the percentage of Ki67-positive cells was increased with IL-22 treatment (Fig. 6G and **S6**). Hence, IL-22 inhibited the hepatocyte apoptosis and increased hepatocyte regeneration. To further evaluate the molecular basis of the liver protective activities of IL-22 on steatohepatitis, mice were fed with high-fat-diet and then treated with IL-22 (2.5 mg/kg) or PBS during high-fat-diet feeding for indicated times (Fig. 6I). We also suggested that IL-22 treatment significantly ameliorated high-fat-diet-induced hepatocellular necrosis, injury, steatosis, ROS accumulation, and mitochondrial dysfunction via activation of AMPK-AKT-mTOR signaling pathways (Fig. 6J and 6K). Consistent with our above observations, real-time PCR assays also revealed that IL-22 distinctly induced lncRNA H19 and glycolytic enzyme (HK-2) expression. Furthermore, we showed that IL-6 and TNF-α in the liver were significantly alleviated by IL-22 treatment (Fig. 6L). Collectively, our findings demonstrated that IL-22 prevented liver injury via activation of mTOR signaling transduction and inhibition of ROS accumulation through mitochondrial function regulation.

**Figure 6.**
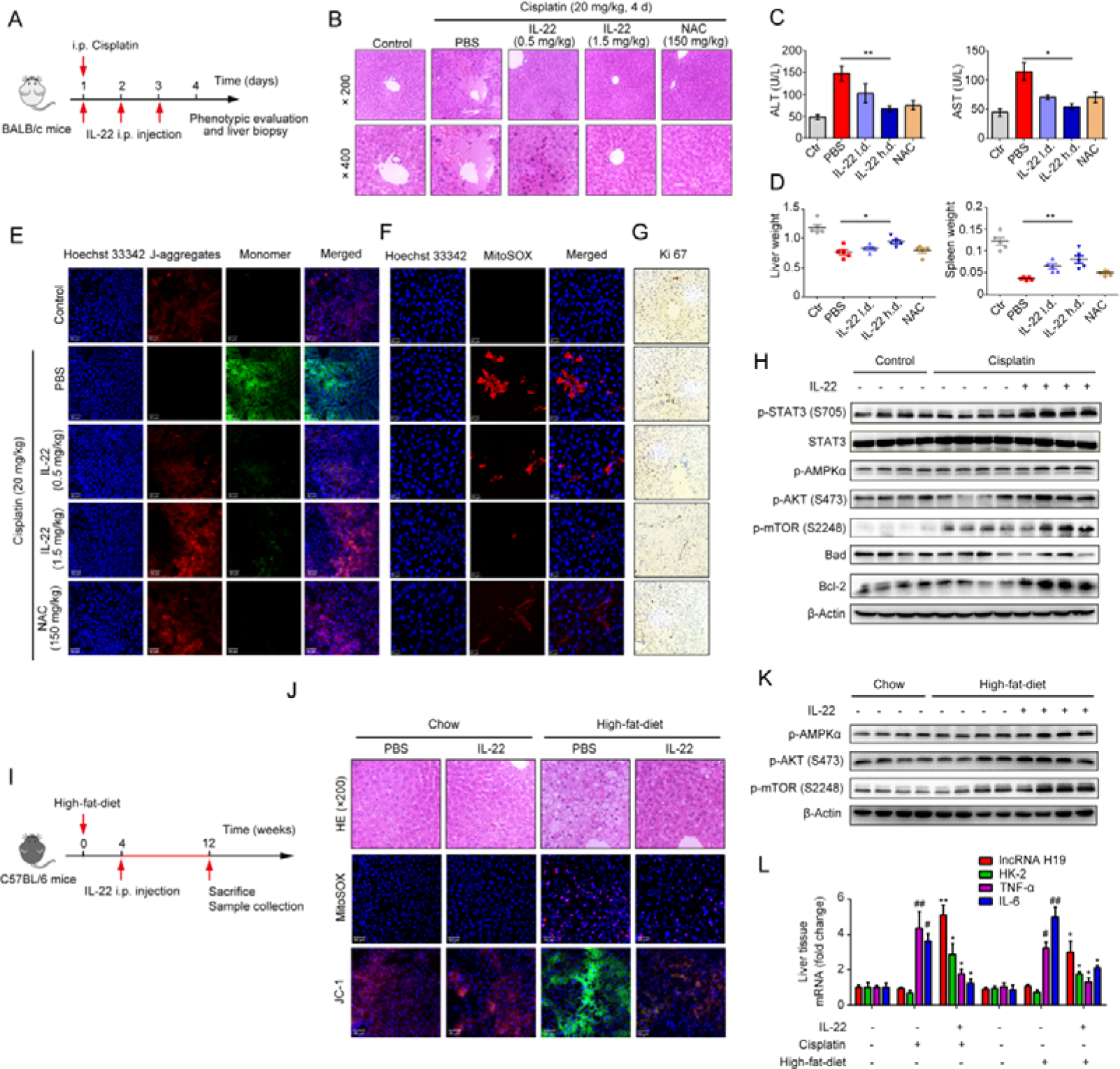
IL-22 attenuates hepatic oxidative stress, mitochondrial dysfunction and damage *in vivo.* (**A**) Schematic diagram of the animal experimental protocols to assess the effects of IL-22 in cisplatin (20 mg/kg) induced liver injury. *N*-acetyl-L-cysteine (NAC) as a positive control group. Representative HE (**B**), MitoSOX (**E**), JC-1 (**F**), and Ki-67 staining (**G**) images of the liver sections were presented. (**C** and **D**) Serum AST levels, serum ALT levels, liver weights and spleen weights were measured. (**H**) Comparison of AMPK/AKT/mTOR activation in liver extracts from IL-22-treated mice versus control subjects was assessed by western blot analysis. (**I**) Schematic diagram of the animal experimental protocols to assess the effects of IL-22 in high-fat-diet fed mice. (**J**) Representative HE, MitoSOX, and JC-1 staining images of the liver sections were presented. (**H**) Liver extracts were subjected to western blot analysis with various antibodies as indicated. (**J**) Western blotting suggested that IL-22 induced AMPK/AKT/mTOR activation in liver extracts from IL-22-treated mice. (**L**) The mRNA expression levels of the indicated genes in the liver (*n* = 3). *P ≤ 0.05 and **P ≤ 0.01 compared with cisplatin-treated and HFD-fed mice. #P ≤ 0.05 and ##P ≤ 0.01 compared with PBS-treated and chow diet fed mice.

## 4. Discussion

Multiple lines of *in vitro* and *in vivo* studies have suggested that IL-22 has significant protective effects against liver injury [13–17]. However, the roles of IL-22 in cell metabolism processes, particularly in hepatocytes and liver, have not been extensively investigated. Previously studies showed that IL-22 ameliorated endoplasmic reticulum (ER) stress and ROS accumulation induced by various stimuli in beta cells via inhibition of oxidative stress associated genes and upregulation of antioxidant gene transcription [13–15, 29]. These findings provided context to our investigation that IL-22 was capable of protecting liver injury by regulating essential metabolic processes. In the current study, we first demonstrated that IL-22 drives a metabolic adaptive reprogramming to maintain mitochondrial fitness, which in turn regulated hepatocellular metabolic processes. Further data indicated that IL-22 was capable of upregulating lncRNA H19, AMPK, mTOR and AKT in hepatocytes, key regulators of cell proliferation and epithelial wound healing, to eliminate dysfunctional mitochondria and opening up a new arena for IL-22 mediated mechanisms of action [28, 30]. Together, these data illustrated that the induction of metabolic reprogramming in hepatocytes could attribute to IL-22-mediated liver protective activities.

Cell metabolism is a promising targetable signaling transduction for treating tissue injury diseases. For instance, increased flux via glycolysis has been suggested as a protection mechanism of cell damage, which is associated with enhancing mitochondrial function through compensating impaired energy generation [22, 31–32]. Moreover, the upregulation of OXPHOS in renal tubular endotheliocyte prevents ischemic reperfusion injury, which may thus heighten the emerging therapeutic approach [33]. To our best knowledge, we first suggested that rather than switching from mitochondrial OXPHOS to glycolysis, IL-22 drove a metabolic reprogramming of hepatocytes via promoting the activities of both. The activation of glycolysis by IL-22 via the facilitation of glycolytic protein expression and glucose uptake indicated that IL-22 could alter the metabolic profiles associated with the hepatocyte injury. These results are consistent with the tissue repair function of Lin28 and also accord with the observed data from the damaged pinnal tissues, where cellular metabolism is needed for tissue function, but if these processes are inhibited, then the tissue protection efficacy is negated [21].

Emerging evidence suggests that mitochondria are the heart of organelles that integrate cell metabolism and cell fate [24–25, 34]. In the present investigation, our findings indicate that IL-22-pretreated hepatocytes maintained mitochondrial fitness with the increased glycolysis and OXPHOS in hepatocytes on injury factors stimulation. Of note, AMPK, a vital energy sensor, regulates cell metabolism to sustain organism homeostasis [35]. Particularly, AMPK also prevents cell or tissue damage by alleviating mitochondrial dysfunction and inhibiting cell apoptosis [36–37]. Consistent with these studies, our results suggesting that IL-22 induces the activation of AMPK signaling demonstrate that IL-22 has more straight-forward actions in sustaining mitochondrial function, which are significant to maintaining hepatocyte respiratory capacity. Additionally, IL-22 signaling pathways through STAT3 may play direct roles in preserving mitochondrial integrity, as these have been demonstrated formerly that the activation of STAT3 signaling exists in mitochondria and has essential roles in the electron transport chains [38]. Moreover, STAT3 has been revealed to potentiate glucose metabolism and accelerate glycolysis. Thus, we can conclude with complete confidence that IL-22 administration alters the metabolic reprogram of hepatocyte *in vitro* and *in vivo*. Future therapeutic and mechanistic researches will concentrate on exploring additional targets of IL-22 that may take part in liver protection.

There is growing recognition that mTOR plays a central role in cell metabolism [39]. In this study, our results illustrate that IL-22 activates mTOR signaling through STAT3, which demonstrate that IL-22 regulates hepatocyte metabolic profiles by means of engaging the controlling of mTOR associated signaling transductions. These data support our aforementioned investigations in multiple ways. Firstly, mTOR signaling transduction is vital to regulate the promotion of OXPHOS and glycolysis in numerous tissues [40–42]. This accords with our observations indicating an exaggerated enhancing glycolysis and OXPHOS in IL-22 pretreated hepatocytes after exposure to injurious stimuli, where the mTOR signaling is activated by IL-22. Secondly, the activation of AMPK-AKT signaling is universally known to induce mTOR signaling, demonstrating a possible mechanism for metabolic reprogramming of hepatocytes by IL-22. Thirdly, it has also been suggested that mTOR signaling activation potentiates glycolytic enzymes expression, likely HK-2, c-Myc [43]. Our findings therefore indicate a stream-lined mechanism for the molecular basis of the liver protective potentials of IL-22, where IL-22 leads to the induction of mTOR activation, causing the upregulation of glycolytic enzymes, as suggested in Fig. 4E, 4F and 4G we observed. There are some issues yet to be addressed, such as if IL-22 also regulates other metabolic processes linked with its hepatoprotective activities via promoting the mTOR associated signaling transductions, and need further research.

Among several regulators of mTOR signaling that have been exported, we first suggest that the expression of lncRNA H19 is remarkably upregulated by IL-22 in hepatocytes and that the IL-22-STAT3-lncRNA H19 signaling pathway is significant for the activation of mTOR and the preservation of mitochondrial fitness during hepatocyte damage. In agreement with previous studies, we show that lncRNA H19 affects the upstream regulators of mTOR signaling, such as AMPK, AKT proteins, thereby activates mTOR and induces the expression of a subset of metabolic reprogramming regulators in hepatocytes [30, 44–46]. Accordingly, we find that lncRNA H19 plays a central role in the inhibition of mitochondrial ROS accumulation and dysfunctional mitochondria elimination in hepatocytes. Further investigations are necessary to explore the underlying mechanism of STAT3-induced lncRNA-H19 expression and fully characterize the *in vivo* functions of lncRNA H19 during liver injury and other liver diseases.

In summary, our findings indicate a crucial role of IL-22 in regulating hepatocellular metabolism through a metabolic reprogramming to maintain mitochondrial fitness and treat liver injury. We put forward that these metabolic regulations by IL-22 are vital to control of hepatocellular function and survive. Lacking their mediators (such as, inhibited lncRNA H19, AMPK, AKT or mTOR activation) can lead to loss of metabolic reprogramming capacities of IL-22 and abnormal mitochondrial fitness, as shown in damaged hepatocytes from IL-22-pretreated cells, which have restrained related signaling transductions. Notably, the liver injury by ROS accumulation owing to dysfunctional mitochondria in animal models was significantly ameliorated by IL-22 via activation of mTOR signaling. Our data are particularly implication to the area for several reasons. Firstly, the controlling of mTOR, a conservative metabolic regulator, by the IL-22-STAT3-lncRNA H19-AMPK-AKT-mTOR axis has not been previously demonstrated. Secondly, therapeutic targeting of these metabolic processes in hepatocytes therefore can be directly favorable to the prevention or treatment of liver injury diseases. Finally, with recombinant IL-22 proteins at present being explored as potential therapeutic strategies in human diseases as severe alcoholic hepatitis, graft-versus-host disease and inflammatory bowel disease, our research may have momentous clinical significances [18–19, 37].

## 3. Materials and Methods

### Reagents and antibodies

Recombinant IL-22 proteins were obtained from Novoprotein, China; oligomycin, cyanide p-trifluoromethoxyphenyl-hydrazone (FCCP) and rotenone were purchased from Seahorse Biosciences, USA; 2-Deoxy-D-glucose, glucose, palmitic acid (Palm.), rapamycin were purchased from Sigma-Aldrich, USA; dorsomorphin, LY294002, cisplatin (Cisp.), 5,5′,6,6′-tetrachloro-1,1′,3,3′-tetraethylbenzimidazolylcarbocyanine iodide (JC-1), TRNzol reagent, MitoTracker Green, SYBR Green qPCR mix, MMLV reverse transcriptase were provided by Beyotime Biotechnology, China; ethanol (Eth.), carbon tetrachloride (CCl_4_) were obtained from Sinopharm, China; MitoTracker Red, Hoechst33342, PKH26, MitoSOX were purchased from Invitrogen, USA. Collagenase IV was obtained from Yeasen, China. Percoll was purchased from GE, USA. Antibodies targeting AKT, p-AKT (S473), mTOR, p-mTOR (S2248), p-STAT3 (Y705), p-PI3K (Tyr199), p-p70S6K, HK-2, c-Myc, Bad, Bcl-2 were obtained from Cell Signaling Technology, USA; Antibodies for Glu1, AMPK, p-AMPKα, IL-22R1, β-Actin and GAPDH were provided by Abcam, USA.

### Hepatocyte culture and stimulation

Hepatocytes were isolated from male C57BL/6 mice livers by a nonrecirculating, retrograde perfusion with 0.08% type IV Collagenase and then were purified and collected by the Percoll solution (40%). Human hepatocyte cell line L02 was obtained from Cell Bank of Chinese Academy of Science. Hepatocytes were incubated in standard culture medium containing 10% fetal bovine serum (FBS), Dulbecco’s modified Eagle’s medium (DMEM) and 1% streptomycin-penicillin, and were grown in a 37°C atmosphere with 5% CO_2_. We incubated hepatocytes with IL-22 (0.5 μg/mL) for 0.5 h, then 200 mM ethanol or 5 μg/mL cisplatin or 0.25 mM palmitic acid or 10 mM CCl_4_ for 24 h. In some cases, hepatocytes were cultured in the presence of dorsomorphin (5 μM), LY294002 (20 μM) or rapamycin (50 nM) for indicated times.

### Seahorse experiments

We tested hepatocytes using a Seahorse XF96 Extracellular Flux Analyzer, for changes in the oxygen consumption rate (OCR, pmol O_2_/min) and extracellular acid rate (ECAR, mpH/min) as a measure of OXPHOS and glycolysis respectively. Briefly, hepatocytes were planted overnight on 96-well polystyrene Seahorse plates. Before starting the experiments, we washed and cultured hepatocytes with seahorse assay mediums containing 2 mM glutamine, 1 mM pyruvate and 10 mM glucose (or without glucose for analyzing ECAR) in a 37°C atmosphere without CO_2_ for 30 min. Experiment results were measured at the indicated time points and after the injection of the under-mentioned inhibitors at optimum concentrations of oligomycin (1.0 µM), FCCP (1.0 µM), rotenone/antimycin A (0.5 μM), glucose (10 mM) and 2-DG (50 mM).

### Flow cytometry

Hepatocytes were cultured and stimulated as mentioned above. MitoTracker Red (mitochondrial membrane potential), MitoSOX (mitochondrial ROS) and MitoTracker Green (total mitochondrial mass) staining were performed in accordance with the manufacturer’s protocols and previous studies [14–15]. Results were obtained by a Beckman Coulter Flow Cytometer (BD Biosciences) and analyzed with CytExpert software.

### Gene knockdown

Small interfering RNA (siRNA) was provided by RiboBio (Guangzhou, China). We prepared transfection cocktails by mixing siRNA, (100 pmol) with Lipofectamine RNAiMAX via gentle pipetting and cultured them at room temperature for 30 min. For siRNA gene silencing, we removed the standard culture mediums and then used the transfection cocktails to incubate the hepatocytes (1×10^6^). After 6 hours incubation in an incubator, the transfection cocktails were gently changed by fresh culture mediums. Forty-eight hours after transfection, hepatocytes were treated with IL-22 for the following experiments.

### Immunofluorescence

Hepatocytes grown in microscopy chambers were fixed with 4% paraformaldehyde, permeabilized with 0.1% Triton X-100, and blocked with 10% bovine serum albumin (BSA) for 60 min. These hepatocytes were then stained with Alexa-488 conjugated anti-GLUT1 antibody at 4°C overnight, washed three times, and stained with PKH26. After washing, hepatocytes were mounted on slides with anti-fade mounting medium containing Hoechst 33342. Pictures were acquired on a confocal microscopy (Zeiss-710, Germany).

### RNA-Seq analysis

Total RNA was extracted from cell samples (1×10^7^) using TRNzol reagent (Beyotime Biotechnology, China). Sample sequencing was performed on an Illumina HiSeq sequencing system at 50-base read length.

### Immunoblot analysis

The immunoblot analysis was performed as previously stated [13–15]. Briefly, we subjected protein samples to sodium dodecyl sulfate (SDS)-polyacrylamide gel electrophoresis and transferred the separated proteins to polyvinylidene (PVDF) membranes. Subsequently, these membranes were blocked with BSA for 2 hours and then incubated with the primary antibodies at 4°C for 12 hours. After washing four times, these membranes were subjected to horseradish peroxidase (HRP)-conjugated secondary antibody and detected using an enhanced chemiluminescence instrument (Pierce, USA). *In situ* hybridization was performed using the miRCURY LNA microRNA *in situ* hybridization kit (Exiqon, Denmark) in accordance with the manufacturer’s manual. The images were captured using the confocal microscopy.

### Glucose uptake assay

The concentrations of glucose in hepatocyte culture supernatants were detected using the glucose uptake assay kit as the manufacturer’s instructions (GAHK20, Sigma). Values were normalized to cell number or tissue mass, as appropriate

### Real-time PCR

Total RNA was obtained from cell samples (1×10^6^) by TRNzol reagent and was transcribed to cDNA by MMLV reverse transcriptase kit. Then the expression levels of mRNA were measured on a BioRad real-time PCR instrument using SYBR green qPCR-mix kit and normalized to GAPDH.

### Mice and histological assay

C57BL/6 and BALB/c mice were provided by Slaccas Experimental Animal Co. (Shanghai, China) and were housed in specific pathogen free (SPF) facilities at 22°C with 12 h light/dark cycles. For the drug-induced liver injury model, mice were injected intraperitoneally with cisplatin (20 mg/kg) or saline control. The C57BL/6 mice were fed with normal chow diets or high-fat diets for the indicated times to induce steatohepatitis. The animal experiments were performed following the protocols and procedures approved by the Institutional Animal Care and Use Ethics Committee (IACUE) at Fudan University. ROS, mitochondrial membrane potential, immunohistochemical and histological staining of liver sections were performed as mentioned previously [14].

### Statistical Analysis

Results were expressed as means ± standard deviations (SD) unless specified differently. Statistical analyses of experimental results were evaluated using GraphPad Prism 5.0 (LaJolla, USA). Differences were analyzed using one-way of analysis (ANOVA) or Student’s t-test. Statistical significance was shown as ***P < 0.001, **P < 0.01 or *P < 0.05.

## List of Abbreviations

IL-22: interleukin 22
OXPHOS: oxidative phosphorylation
mTOR: mammalian target of rapamycin
ROS: reactive oxygen species
IL-10: interleukin-10
ILCs: innate lymphoid cells
OCR: oxygen consumption rate
ECAR: extracellular acid rate
FCCP: cyanide p-trifluoromethoxyphenyl-hydrazone
CCl_4_: carbon tetrachloride
FBS: fetal bovine serum
BSA: bovine serum albumin
IL-22R1: IL-22 receptor
MRC: maximal respiratory capacity
KEGG: Kyoto Encyclopedia of Genes and Genomes analysis
SSC: side scatter signal
AMPK: AMP-activated protein kinase
ALT: alanine aminotransferase
AST: aspartate amino-transferase
ER: endoplasmic reticulum

